# Resistance to BRAF inhibitors drives melanoma sensitivity to Chk1 inhibition

**DOI:** 10.1101/2025.01.14.632956

**Authors:** Danielle G. Carvalho, Juliana C. N. Kenski, Daniel A. Moreira, Matheus A. Rajão, Oscar Krijgsman, Carolina Furtado, Mariana Boroni, João P. B. Viola, Daniel S Peeper, Patricia Abrao Possik

**Author notes:** Contributed equally (Danielle G. Carvalho, Juliana C. N. Kenski). Lead Contact: Patricia A. Possik.

## Abstract

BRAF inhibitor-resistant melanomas (BRAFiR) acquire (epi)genetic and functional alterations that enable them to evade alternative treatments. Identifying these alterations is critical to advancing treatment strategies. Here, we explored the effect of Chk1 inhibition (Chk1i) on BRAFiR cells, revealing higher sensitivity compared to treatment-naïve cells both *in vitro* and *in vivo*. Using FUCCI-labeling and time-lapse microscopy, we show that S phase progression is required for Chk1i-induced cytotoxicity in BRAFiR cells, but not in treatment-naïve cells. Replication stress markers, including reduced BrdU incorporation and increased phospho-RPA and γH2AX, were exclusive to BRAFiR cells exposed to Chk1i. Untreated BRAFiR cells exhibited upregulated DNA replication genes, reduced progressing forks and increased origin firing, suggesting intrinsic replication changes. MAPK pathway reactivation in treatment-naïve cells mimicked BRAFiR traits, increasing sensitivity to Chk1i. These findings indicate that Chk1i exploits elevated replication stress specifically in BRAFiR, highlighting its therapeutic potential in overcoming MAPK inhibitor resistance in melanoma.

## INTRODUCTION

Cutaneous melanoma is a very aggressive form of cancer, accounting for most deaths associated with skin cancer worldwide^1^. Activating mutations in *BRAF*, most commonly *BRAF*^V600E^, serve as genetic drivers in about half of cutaneous melanoma patients, leading to constitutive activation of the MAPK pathway^2^. This activation results in hyper-phosphorylation of retinoblastoma protein (Rb), triggering the liberation of E2F transcription factors and promoting progression through G1 and S phases of the cell cycle^3^. The high frequency of *BRAF*^V600^ mutations led to the development and approval of mutant BRAF and MEK inhibitors for the treatment of advanced cutaneous melanoma^4–11^. However, despite significant clinical benefits, patients typically develop therapeutic resistance within months^7,11–16^. MAPK pathway reactivation is the primary mechanism of resistance and is driven by different (epi)genetic alterations, including in tyrosine kinase receptors, BRAF, MEK, and NRAS^17–38^. The high prevalence of therapeutic resistance poses a significant challenge for durable clinical benefit of patients treated with MAPK pathway inhibitors. Therefore, increasing our understanding of the common mechanisms of resistance is crucial to identify new therapeutic targets.

The DNA damage response (DDR) is an important signaling network responsible for detecting and repairing chemical or structural alterations in the DNA, protecting the cell against the deleterious effects of such DNA lesions. Among other functions, the DDR pathway regulates cell cycle progression, DNA damage repair and apoptosis^39^. The therapeutic potential of DDR inhibitors has been widely explored, since tumor cells often have increased DNA replication rates and are more dependent on the activity of DDR factors to control DNA damage levels, rendering them more susceptible to their inhibition^40^.

Checkpoint kinase 1 (Chk1) plays an essential role in DDR signaling. It is the key factor responsible for preventing or delaying S phase progression^41–43^ and inhibiting premature entry into mitosis in the presence of DNA damage^44,45^. Pharmacological inhibition of Chk1 increases replication stress levels, promotes accumulation of DNA damage and induces cell death in several tumor cell lines^46–55^.

Preclinical *in vitro* and *in vivo* studies on melanoma have demonstrated that response to Chk1 inhibitor was associated with high basal levels of replication stress and DNA damage^56,57^. Furthermore, in the context of therapeutic resistance, previous research suggested that melanoma cells resistant to MAPK inhibitors were also sensitive to Chk1 inhibitors, with this sensitivity being even more pronounced in specific resistant cells^52,55,58,59^. However, the mechanism by which resistant cells respond better to Chk1 inhibition remains unknown and a potential biomarker is lacking. Here, we evaluated and characterized the effects of Chk1 inhibition in melanoma cell lines resistant to BRAF^V600^ inhibitor treatment and in their respective treatment-naïve parental lines.

## RESULTS

### BRAF inhibitor-resistant melanoma cells are hypersensitive to GDC-0575

To comparatively evaluate the effect of Chk1 targeting in the context of BRAF inhibitor resistance, we performed dose response curves with the highly selective Chk1 inhibitor GDC-0575, using five treatment-naïve (or parental, P) melanoma cell lines and their corresponding BRAF inhibitor-resistant derivatives (BRAFiR, R)^29^. Three BRAFiR cell lines (888mel, M026.X1.CL and A375) demonstrated increased loss of viability (as judged by a significantly lower IC50) following GDC-0575 exposure compared to their treatment-naïve counterparts (**Figure 1A**). This included a cell line derived from a patient who acquired resistance to vemurafenib after initially responding to treatment (PDX cell line M026R.X1.CL)^36^. Among the five pairs of tested treatment-naïve and BRAFi-resistant counterparts (**Figure 1**; **Supplementary Figure 1A and B**), four pairs demonstrated hypersensitivity of the resistant cells to Chk1 inhibition. This indicates that this phenomenon is quite common, although not universal, as only one cell line pair tested did not exhibit this response (D10). Increased sensitivity to Chk1 inhibition was also observed in a melanoma cell line with acquired resistance to combined BRAF and MEK inhibitors **(Supplementary Figure 1A and B)**, suggesting that this vulnerability is not limited to BRAF inhibition, but also applies to inhibition of other kinases of the MAPK pathway.

**Figure 1:**
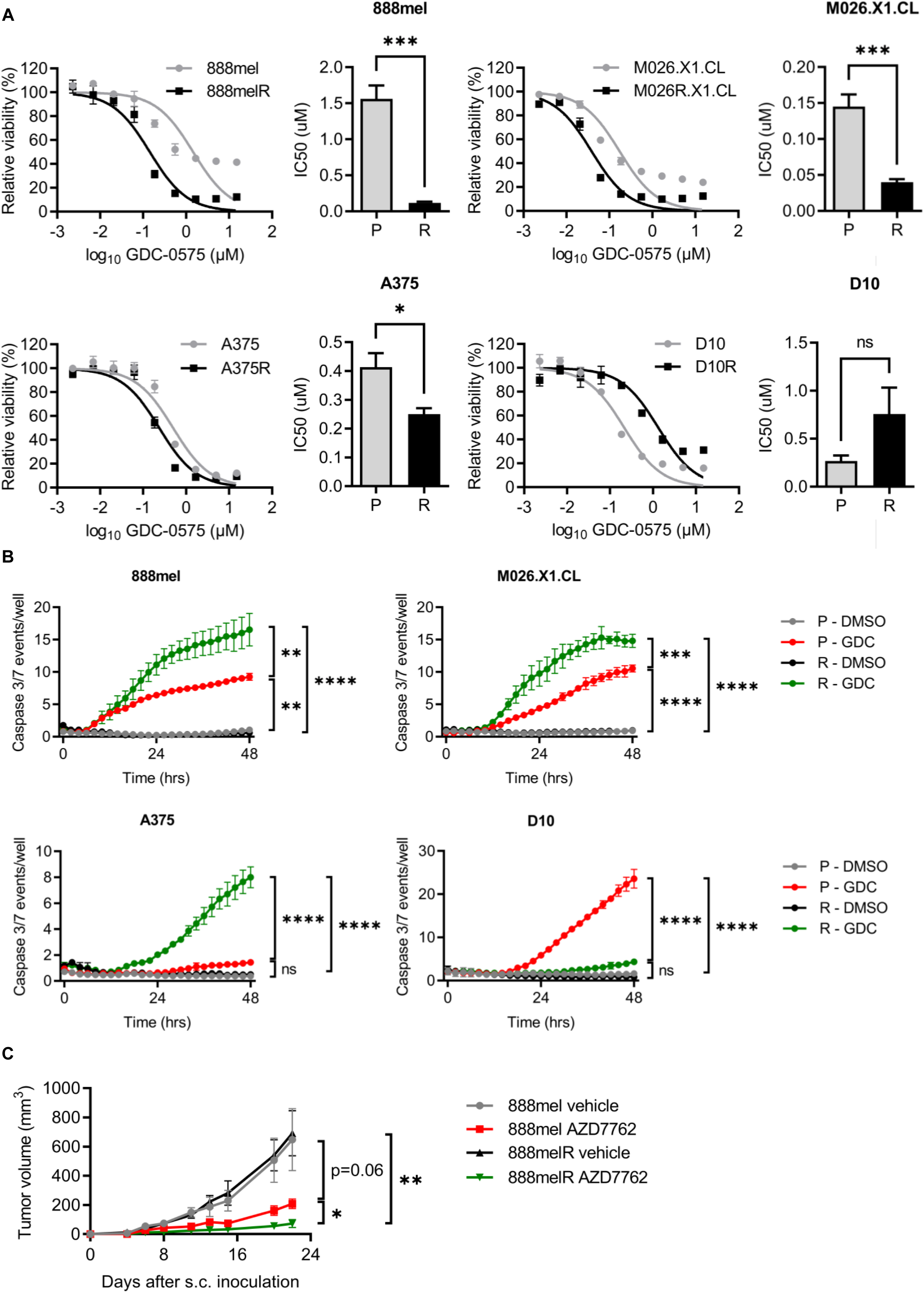
BRAF inhibitor-resistant melanoma cells are hypersensitive to GDC-0575. A) Treatment-naïve parental (P) and BRAF inhibitor-resistant (R) melanoma cells were treated with increasing concentrations of GDC-0575 for 72 hours. Cell viability was determined by CellTiter Blue. The percentage of viable cells are depicted as mean (+/- SD). IC50 values were estimated from the curves and represented as mean (+/- SEM). n = 3 biological replicates with 3 technical replicates each. Unpaired Student#x2019;s t test or Mann Whitney (+/- SEM): * p < 0.05; *** p ≤ 0.001; ns: not significant. B) Cells were exposed to GDC-0575 (GDC, 0.5µM for 888mel/888melR, A375/A375R and D10/D10R and 0.1µM for M026.X1.CL/M026R.X1.CL) or DMSO in the presence of cleaved-caspase dye for 48 hours and monitored using time-lapse microscopy. Cell death was analyzed at each time-point by quantification of green object count normalized by the percentage of phase object confluence. Each time frame is indicated as mean (+/- SD). n = 3 biological replicates with 3 technical replicates each. One representative experiment is shown. Two-way ANOVA. * p < 0.05; *** p < 0.001; **** p < 0.0001. C) 888mel and 888melR cells were inoculated s.c. into NSG mice. Vehicle or AZD7762 treatment (25mg/kg) started at day 1 after tumor inoculation and was performed 3x week. n=4 mice per group. Unpaired Student’s T Test. * p < 0.05.** p < 0.01.

To investigate whether the observed differences in cell viability were indicative of increased cell death, we used microscopy-assisted live cell imaging and temporal measurements of Caspase 3/7 cleavage. Induction of apoptosis was more pronounced in the same three BRAFiR cell lines compared to their parental counterparts (**Figure 1B**, **Supplementary Figure 2A**). In line with what we observed above, the only exception was D10R cell line, which showed lower levels of cell death compared to D10 parental cells. Together, *in vitro* viability and cell death experiments consistently demonstrated that the majority of MAPK inhibitor-resistant melanoma cell lines were more sensitive to Chk1 inhibition than treatment-naïve cell lines.

To confirm these results *in vivo*, we inoculated 888mel and 888melR cells subcutaneously and xenografts were treated with the Chk1 inhibitor AZD7762 or vehicle control. AZD7762 treatment resulted in a significantly higher tumor growth inhibition in BRAFi-resistant cells (**Figure 1C**), which therefore resulted in increased survival (**Supplementary Figure 2B**). Overall, these results demonstrate that 4 out of 5 melanoma cell lines, including a PDX cell line, with acquired resistance to MAPK pathway inhibitors become hypersensitive to Chk1 inhibition both *in vitro* and *in vivo*.

### Chk1 inhibition causes cell cycle failure in BRAF inhibitor-resistant melanoma cells

Sensitivity to Chk1 inhibitors can be related to higher levels of endogenous replication stress, which accelerates premature entry into mitosis^47^. Considering that Chk1 regulates origin firing during S phase and mitotic entry during G2^42,44,45^, we investigated whether alterations in cell cycle checkpoints were associated with BRAF inhibitor resistance and Chk1 inhibitor sensitivity. We used the Fluorescence Ubiquitination-based Cell Cycle Indicator (FUCCI) system^60^, which allows studying cell cycle dynamics by differential labeling of the nuclei of cells in G1 phase (red), early S phase (yellow), and late S, G2 phase or mitosis (green) (**Figure 2A**, **Supplementary Figure 3A**). Firstly, we estimated cell cycle duration of individual FUCCI-labeled cells under normal conditions and observed that BRAFiR and treatment-naïve melanoma cells display similar cell cycling profiles (**Supplementary Figure 3B, C**).

**Figure 2:**
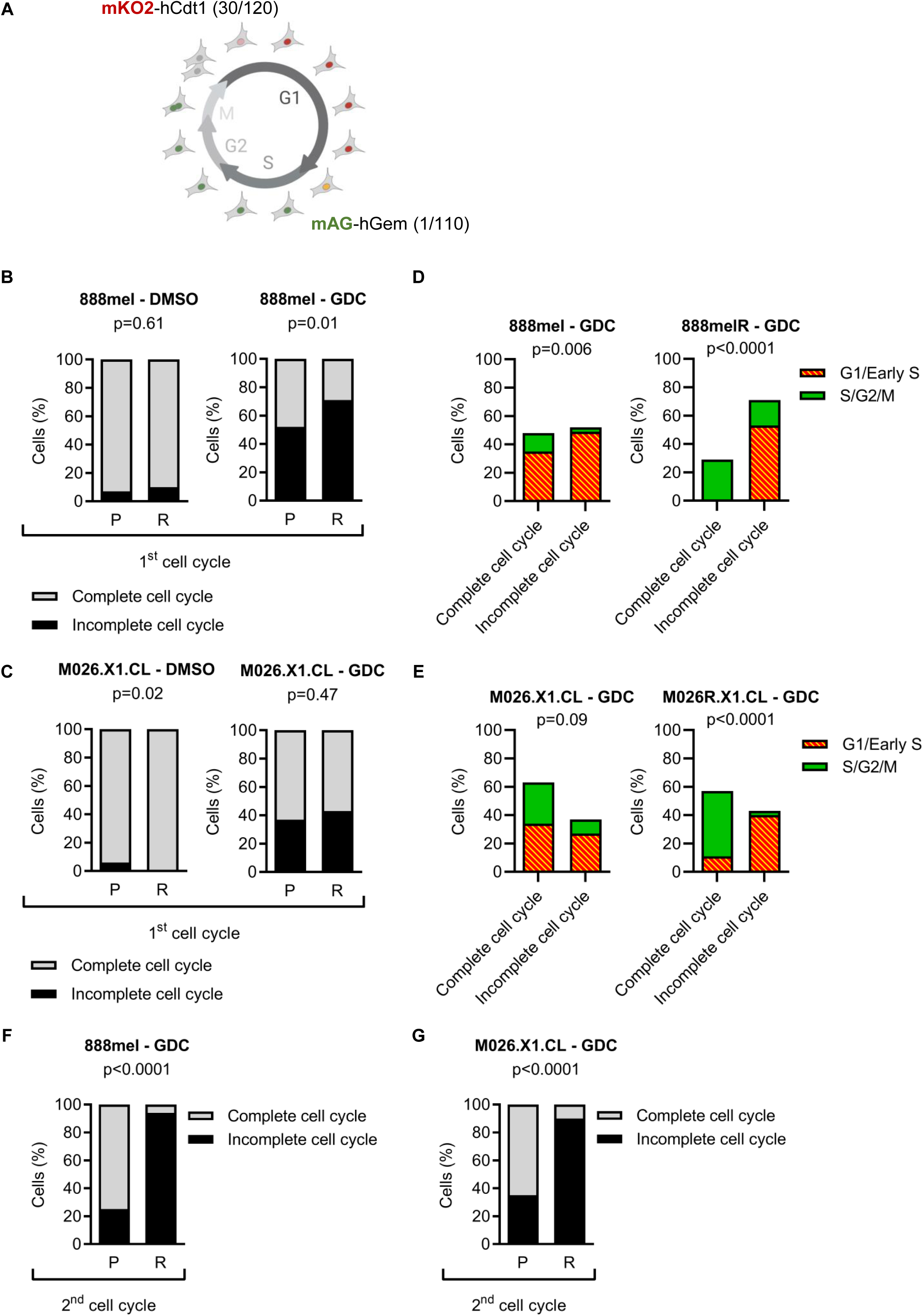
Chk1 inhibition causes cell cycle failure in BRAF inhibitor-resistant melanoma cells. FUCCI-labeled treatment-naïve parental (P) and BRAF inhibitor-resistant (R) melanoma cells were treated with GDC-0575 (0.3µM for 888mel/888melR and 0.1µM for M026.X1.CL/M026R.X1.CL) or DMSO and followed by time-lapse microscopy for 90 hours. A) Schematic representation of the FUCCI system. B-C) Percentage of cells that completed or failed to complete one cycle of cell division. The first cycle was analyzed in 21-34 cells for each condition. D-E) The percentage of cells that completed or failed to complete the cell cycle after GDC-0575 treatment were classified according to cell cycle phase at treatment start. A total of 30-34 cells were analyzed for each cell line. F-G) Percentage of daughter cells that completed or failed to complete the second round of cell division. The second cycle was analyzed in 16-21 daughter cells treated with GDC-0575. Chi-squared and Fisher’s exact test. p values are indicated.

Next, we analyzed the effect of Chk1 inhibition on cell fate. For this, we followed individual cells exposed to GDC-0575 or vehicle control over time. For 888mel, a BRAFi-sensitive cell line, approximately 50% of the cells were able to complete cytokinesis upon Chk1 inhibition, which decreased to 30% on 888melR cells (**Figure 2B**). For M026.X1.CL cell lines, approximately 60% of both BRAFi-sensitive and resistant cells were able to complete the cell cycle (**Figure 2C**). The remaining cells failed to divide. Among those, we observed three main phenotypes: cells with a rounded morphology and loss of plate adherence; cells that progressed through cell cycle phases but, instead of dividing, changed from late S/G2/M to G1 without going through cytokinesis; and cells that failed to complete the cell cycle in up to 90 hours (**Supplementary Figure 4A**). The most common phenotype within the latter was the acquisition of a rounded morphology (**Supplementary Figure 4B and 4C**) and almost all cells were in late S or in G2/M by 90 hours (**Supplementary Figure 4D and 4E**). Although no cell death or cell viability marker was used in this experiment, the detachment of cells with rounded morphology likely represented cell death induced by Chk1 inhibition, in line with the observations described above (**Figure 1**).

Time-lapse microscopy enabled the monitoring of individual cells as they underwent one complete round of cell division, followed by the tracking of the resulting daughter cells through a subsequent round of division. Although more than 90% of the untreated parental and BRAFiR cells successfully completed the first round of cell division, upon exposure to Chk1 inhibitor a higher frequency of BRAFiR cells were unable to complete the cell cycle when compared to the parental cells (52% in 888mel vs. 71% in 888melR; 37% in M026.X1.CL vs. 43% in M026R.X1.CL) (**Figure 2B** and **2C**). Interestingly, nearly all BRAFiR cells (888melR and M026R.X1.CL) capable of completing cell division were initially at late S, G2, or mitotic phases upon exposure to GDC-0575 treatment. Conversely, those initially in G1 or early S phases failed to complete cell division (**Figure 2D** and **2E**). The same phenotype was not observed for treatment-naïve cells (**Figure 2D** and **2E**). Consistently, monitoring daughter cells for a second round of cell division revealed an even more pronounced effect of GDC-0575 (**Figure 2F** and **2G**). These observations indicate that progression through S phase is critical for the sensitivity to Chk1 inhibition of BRAFi-resistant melanoma cells.

### Chk1 inhibition increases replication stress in BRAF-inhibitor resistant melanoma cells

To further characterize the effect of Chk1 inhibition in cell cycle progression, we analyzed BrdU incorporation upon exposure to GDC-0575. The data revealed a prevalence of 888melR and A375R cells that, upon Chk1 inhibition, exhibited reduced or no incorporation of BrdU. This occurred without significantly altering the total number of cells in S phase, indicating that Chk1 inhibition promotes a delay or a blockade in DNA replication (**Figure 3A-B**, **Supplementary Figure 5A-B**). M026R.X1.CL cells, on the other hand, accumulated in the beginning of S phase, indicating a potential arrest in cell cycle progression, but this was also observed for M026.X1.CL cells, in which the S phase distribution profile was not altered by Chk1 inhibition (**Supplementary Figure 5A-B**). Both phenotypes indicate Chk1-induced replication stress in BRAFiR melanoma cells. Consistently, upon Chk1 inhibition, we noted a more pronounced increase in phospho-RPA32, a marker of single-stranded DNA, and ɣH2AX, a marker of DNA double strand breaks in BRAFiR melanoma cells that are hypersensitive to Chk1 inhibition (**Figure 3C**).

**Figure 3:**
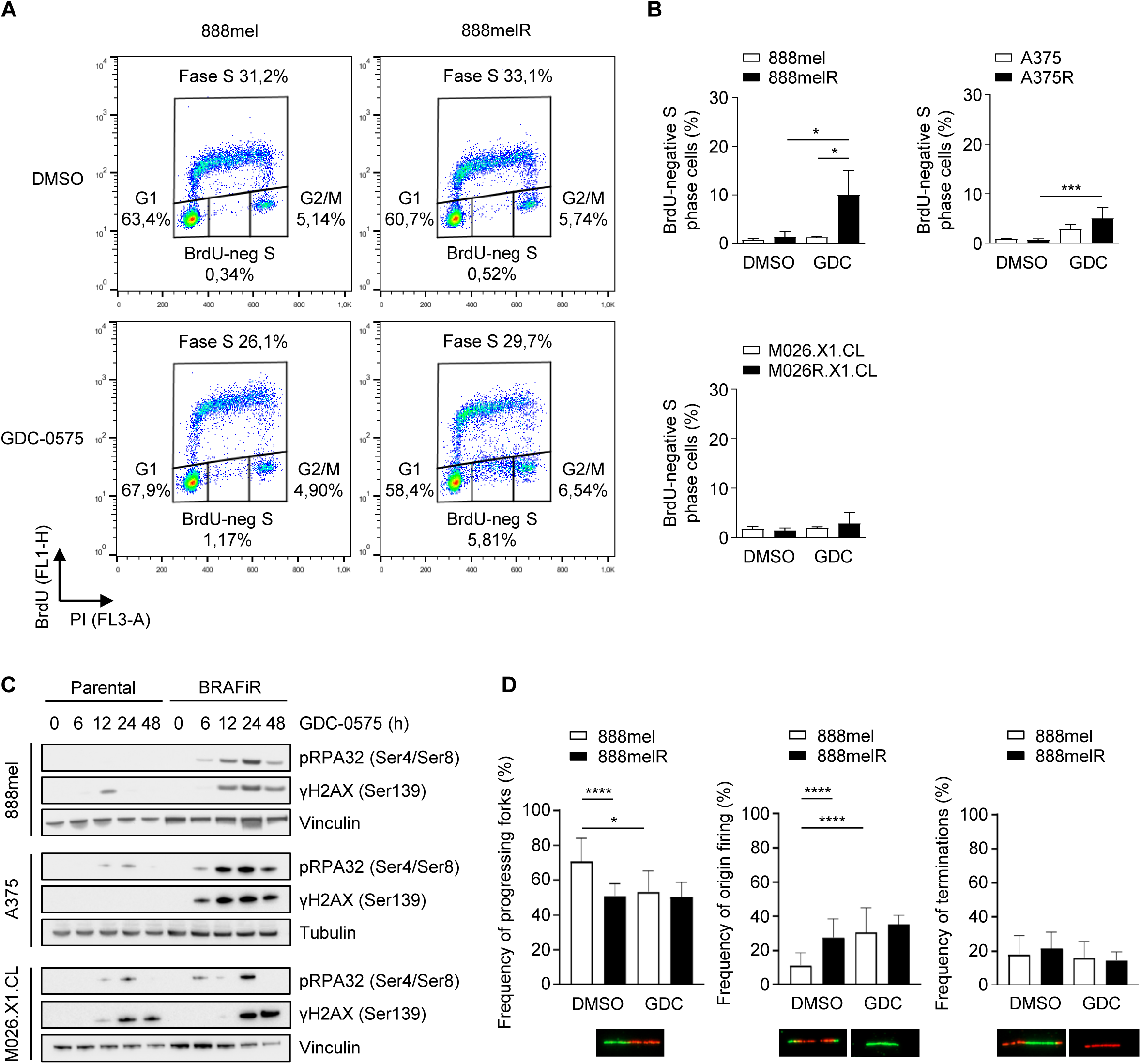
Chk1 inhibition increases replication stress in BRAF-inhibitor resistant melanoma cells. Treatment-naïve parental (P) and BRAF inhibitor-resistant (R) melanoma cells were exposed to GDC-0575 (0.3µM for 888mel/888melR and A375/A375R or 0.1uM for M026.X1.CL/M026R.X1.CL) or DMSO. A) Cells were pulsed with BrdU and analyzed six hours after treatment. n = 3 biological replicates. B) Quantification of BrdU-negative S phase cells from A. C) Western blot analysis at different time points of Chk1 inhibitor treatment. Vinculin and tubulin served as loading controls. n = 3. D) Quantification of the frequency of progressing forks, origin firing and termination in the presence and absence of Chk1 inhibitor. n = 2 biological replicates, and only one is represented. Ordinary one-way ANOVA or Kruskal Wallis. * p < 0.05; *** p < 0.001; **** p < 0.0001. Only significant comparisons are shown.

To further examine the dynamics of S phase progression during Chk1 inhibition, we performed DNA fiber staining. This assay uses pulses of CldU and IdU nucleotide analogs to label nascent DNA, allowing to track the dynamics of DNA synthesis. Chk1 inhibition disrupted DNA replication in both parental and BRAFiR cells, as indicated by a reduction in replication fork speed following treatment (**Supplementary Figure 6A-B**). However, important differences between parental and BRAFiR cells were revealed when analyzing the replication events. In response to Chk1 inhibition, a decrease of progressing forks and an increase in origin firing was only observed in the parental cells. This indicates that DNA stalling and activation of dormant licensed origins, which are predominant characteristics of replication stress, are also occurring in Chk1 inhibitor-treated parental cells^61–63^ (**Figure 3D**). However, we observed that BRAFiR cells already exhibited a lower frequency of progressing forks and a higher frequency of origin firing under normal conditions, suggesting an intrinsic alteration in their replication capacity (**Figure 3D**). Altogether, these results demonstrate that DNA replication is significantly disrupted by Chk1 inhibition in BRAFi-resistant cells, which have higher levels of replication stress than treatment-naïve cells, even in the absence of treatment.

### BRAF inhibitor-resistant cells exhibit increased expression of genes associated with DNA replication pathway

To further explore the intrinsic alterations of DNA replication machinery observed in BRAFiR cells, RNA sequencing was performed in both BRAFi-sensitive and -resistant cells upon CHK1 inhibition. A detailed analysis of pathways related to DNA synthesis and DNA damage response revealed significant suppression of the DNA Replication pathway in both parental and BRAFi-resistant cell lines (**Supplementary Figure 7**). Moreover, closer examination of this pathway indicated that, in the absence of Chk1 inhibitor treatment, 888melR cells exhibit higher basal expression of DNA replication factors such *RFC3*, *PCNA*, *RNASEH2B*, *FEN1*, *POLA2*, and *RFC4*, in comparison to 888mel cells. These genes encode proteins associated with the formation of the replisome and are therefore essential for the DNA replication fork progression. Upon Chk1 inhibition, most of these genes maintain high expression levels (**Figure 4A**). Genes associated with S phase entry, such as *MYC*, *E2F1* and *CCND1,* were also significantly upregulated in untreated BRAFiR cells (**Figure 4B**). These findings indicate that 888melR cells harbor intrinsic alterations capable of modulating the expression of genes that are essential for cell cycle progression.

**Figure 4:**
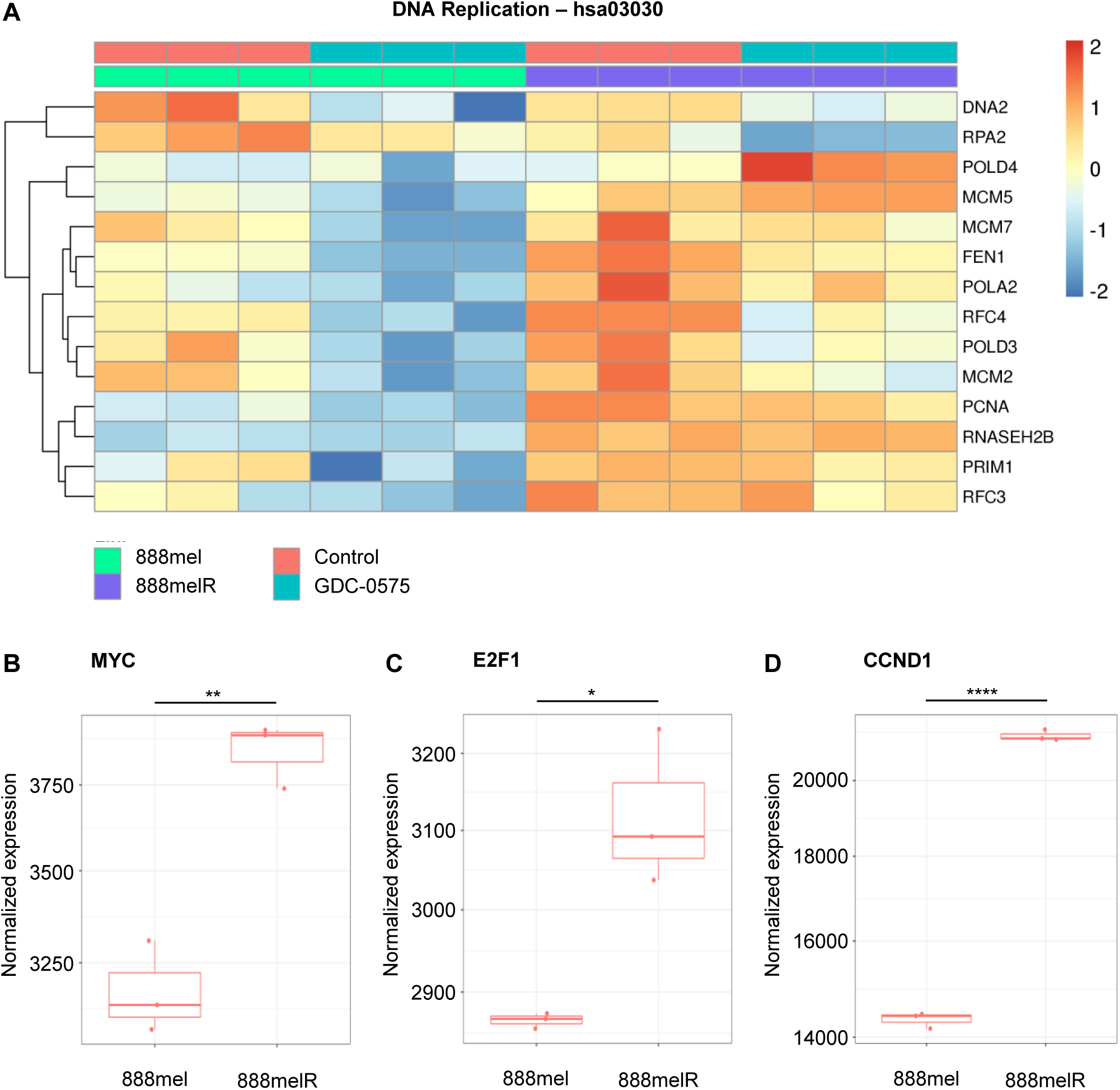
BRAF inhibitor-resistant cells exhibit increased expression of genes associated with DNA replication pathway. RNA sequencing of 888mel and 888melR cells untreated or treated with 0.3 µM GDC-0575 for six hours. n = 3 biological replicates. A) Heatmap depicting genes that belong to the DNA replication pathway accordingly to KEGG database (hsa03030) and are differentially expressed in at least one of the comparisons: 888melR control versus 888mel control; 888melR GDC-0575 versus 888melR control; 888mel GDC-0575 versus 888mel control; 888meR GDC-0575 versus 888mel GDC-0575. The color scheme indicates the Z-score scale for each gene. Wald Test. p-adj < 0.05. Quantification of *MYC* (B), *E2F1* (C) and *CCND1* (D) normalized read counts in untreated samples. Normalization was performed with DESeq2. Unpaired Student’s T Test. * p < 0,05; ** p < 0.005; **** p < 0.00001.

### Activation of the MAPK pathway increases sensitivity to Chk1 inhibition

Our results above have consistently shown that BRAFi-resistant cells not only have intrinsic alterations in DNA replication but are also differentially affected by Chk1-inhibition induced replication stress resulting in hypersensitization. Since reactivation of the MAPK pathway represents the most frequent mechanism of resistance to MAPK inhibitors^64^, we investigated whether this could be responsible for the Chk1 inhibitor hypersensitization phenotype. Indeed, 888melR, A375R and M026R.X1.CL cells consistently showed increased ERK phosphorylation compared to their parental counterparts (**Figure 5A**). To corroborate these findings, we analyzed the expression of ten MAPK-specific genes in the RNA-Sequencing data and quantified the MAPK Pathway Activity Score (MPAS)^82^. This analysis revealed higher baseline MAPK pathway activity in 888melR cells compared to the parental cell line (**Figure 5B**). To validate the role of MAPK pathway activation in Chk1 inhibitor sensitivity, we used the phorbol ester PMA to induce MAPK pathway activation in melanoma cells. As expected, this led to increased ERK phosphorylation (**Figure 5C**) and reduced sensitivity of parental cells to BRAF inhibitor (PLX4720) (**Figure 5D, E**), mimicking what is observed in BRAF inhibitor-resistant cells (**Figure 1A**). Notably, the induction of MAPK pathway signaling sensitized 888mel cells to Chk1 inhibition, as evidenced by a significant decrease in the IC50 values (**Figure 5F, G**). These findings highlight the association between MAPK pathway reactivation and increased sensitivity to Chk1 inhibitors.

**Figure 5:**
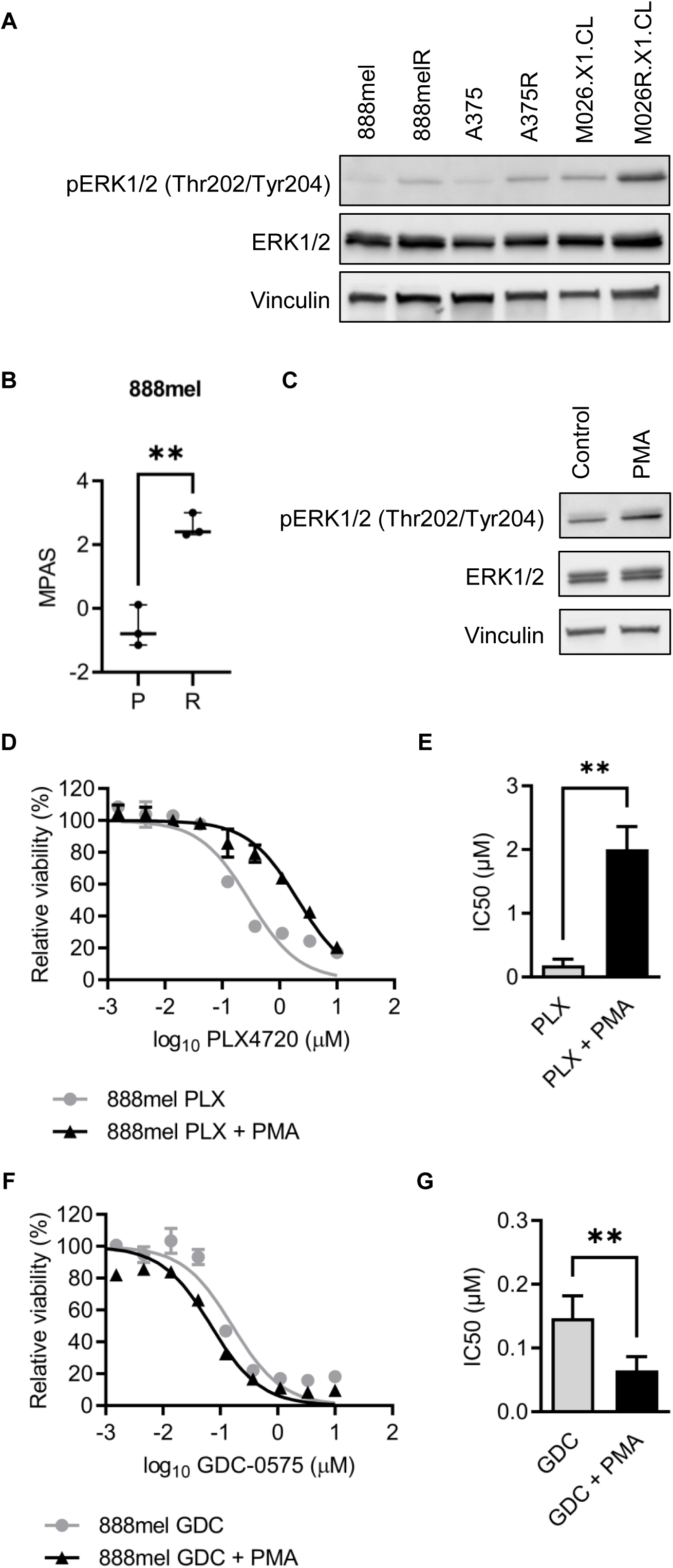
Activation of the MAPK pathway increases sensitivity to Chk1 inhibition. A) Western blot analysis of MAPK pathway components in treatment-naïve parental (P) and BRAF inhibitor-resistant (R) melanoma cell lines. n = 3 biological replicates. B) MAPK Pathway Activity Score (MPAS) for each sample was derived from expression data for 10 MAPK-specific genes^82^. C) Western blot analysis of 888mel cells +/- 10nM PMA for eight hours in DMEM 2% FBS. Vinculin served as loading control. n = 3 biological replicates. D-G) 888mel cells were treated for 72 hours with different concentrations of PLX4720 or GDC-0575 +/- 10nM PMA in DMEM 2% FBS. Cell viability was determined by CellTiter Blue. The percentages of viable cells are depicted as mean (+/- SD) (D and F). IC50 values were estimated from the curves and are represented as mean (+/- SEM) (E and G). n ≥ 3 biological replicates, with 3 technical replicates each. Unpaired Student’s T Test or Mann Whitney. ** p < 0,01.

## DISCUSSION

The development of MAPK inhibitors has significantly improved the treatment of patients with cutaneous melanoma. However, most patients eventually experience disease progression due to treatment resistance^65^. Although extensive research has revealed the main mechanisms of MAPK inhibitors resistance, there is still a need to identify therapeutic strategies to control disease progression in relapsed patients.

Inhibitors of the DNA damage response (DDR) signaling network, such as those targeting ATR and Chk1, have been explored *in vitro*, in preclinical studies and in clinical trials for several tumor types, including melanomas^66,67^. By blocking the DNA damage response, ATR and Chk1 inhibitors limit the ability of cells to repair DNA lesions. Although the principle for using these inhibitors is similar, some effects of Chk1 inhibition are unique, primarily due to the ability to destabilize replication forks during replication stress ^68^. Our previous study has demonstrated that inhibition of Chk1/2 is toxic to melanoma cells, including those resistant to BRAF inhibitors, and that this effect is enhanced in the context of hypoxia^58^. Here, we demonstrate that MAPK inhibitor-resistant melanoma cell lines are more susceptible to killing by Chk1-specific inhibition than their parental counterparts both *in vitro* and *in vivo*. This was observed not only in cell lines resistant to BRAF inhibitors, but also in cells resistant to the BRAF and MEK inhibitors combined. One of these cell line pairs was developed from patients-derived xenografts before and after the acquisition of BRAF inhibitor resistance, reinforcing the clinical relevance of our findings.

Chk1 kinase plays an essential role in regulating cell cycle progression in the response to DNA damage, specifically in the S and G2 checkpoints. Indeed, it has been extensively demonstrated that toxicity induced by Chk1 inhibition is strongly impacted by cell cycle distribution and progression in different tumor models^47,50–55^. In melanoma, Chk1 inhibition promotes the accumulation of cells in the S phase and premature entry into mitosis, leading to an increased rate of aberrant mitoses^47,50,55^. Additionally, previous studies have demonstrated that Chk1 inhibition induces apoptosis in melanoma cells prior to mitotic entry, implying that progression through S phase alone is enough to trigger cell death independently of mitotic catastrophe^53^. Consistent with this, induction of p16 expression blocks entry into S phase and prevents apoptosis induced by Chk1 inhibition in melanoma cells^47^. While our findings align with these observations, we found that progression through the S phase is particularly critical for the effects of Chk1 inhibition in BRAF inhibitor-resistant melanoma cell lines.

By Fucci-labeling of parental and BRAFi-resistant cells, we demonstrate that exposure to Chk1 inhibitor during G1, G1/S transition or early S prevents BRAFi-resistant cells from completing the cell cycle, whereas cells treated in mid or late S, G2, or mitosis were able to complete cell division. Interestingly, this effect was not observed for treatment-naïve cells to the same extent.

In melanoma cell lines, sensitivity to Chk1 inhibition has been correlated with high basal levels of replication stress and DNA damage^47,53^. Although we did not observe intrinsic differences in phosphorylated RPA and γH2AX levels between parental and BRAF inhibitor-resistant cell lines, we showed a reduction of actively replicating cells and an increase of single-stranded DNA and double-stranded DNA breaks upon Chk1 inhibition in BRAF inhibitor-resistant cells, corroborating previous findings^46,47,49–52,54,55^. Moreover, Chk1 inhibition induced an increase of replication origin firing and a decrease of progressing replication forks in the parental cells, but not in the BRAF inhibitor-resistant ones. These results are consistent with previous studies reporting that pharmacological or genetic depletion of Chk1 increases replication origin firing and slows down progression of replication forks^42^. Most importantly, in the absence of any treatment, BRAF inhibitor-resistant cells presented an increase in replication origin firing and a decrease of progressing forks compared to parental cells. This reveals an intrinsic imbalance in the replication machinery of BRAFi-resistant cells, potentially compromising accurate DNA replication and increasing the reliance on Chk1 function in response to DNA damage. The increased expression of transcription factors that control entry into S phase and initiation of DNA replication (*MYC* and *E2F1*) partially explains the increased expression of genes involved in fork progression in BRAF inhibitor-resistant cells.

In addition to the previously proposed mechanisms for sensitivity to Chk1 inhibition cited above, cell cycle regulation also plays an important role in this setting. Activation of the Cyclin A-CDK2 complex, which is crucial for progression through the replication phase of the cell cycle, has been associated with increased DNA damage and subsequent cell death in the presence of Chk1 inhibitor across a wide range of tumor cell lines^42,48,50,69,70^. Although we did not assess the activity of the Cyclin A-CDK2 complex, we identified an increased expression of *CCND1* gene, encoding Cyclin D1, in a BRAF inhibitor-resistant cell line. The Cyclin D-CDK4/6 complex initiates Rb phosphorylation and contributes to the release of E2F transcription factors, an event recently described to be tightly regulated by an intermediate state of Rb-E2F activation dependent on external and internal signals to commit to proliferation and finally enable entry into the S phase^71^. While it is possible that CDK2 activation is involved in the mechanism of hypersensitivity to Chk1 inhibition in our models, further investigation is required to confirm this hypothesis. In lung and colorectal cancer cell lines, silencing of B family polymerases (*POLA1*, *POLE*, and *POLE2*) induces increased sensitivity to Chk1 inhibition^72^. In line with that, our results show that 888melR cells, which are hypersensitive to Chk1 inhibition, exhibit higher expression of *POLA2* compared to its less sensitive parental cell line. Loss of p53 activity has also been proposed as a mechanism for Chk1 inhibitor sensitivity, as p53-deficient tumors lose control of the G1/S transition checkpoint and become more vulnerable to agents that block the S and G2 checkpoints, such as Chk1 inhibitors^73^. However, both parental and BRAFi-resistant melanoma cell lines used here do not have genetic alterations in *TP53* (data not shown), suggesting that, in our models, sensitivity to Chk1 inhibition is independent of p53 loss.

Previous data indicates that MAPK pathway activation is associated with Chk1 activity. Lee and colleagues demonstrated that MAPK signaling positively regulates *CHEK1* transcription and that Chk1 functions as an inhibitor of RAS, preventing its hyperactivation^51^. This regulatory interplay between RAS and Chk1 allows tumor cells with constitutive activation of the MAPK pathway to proliferate without excessive DNA damage accumulation. Gilad and collaborators also contributed to understanding the relationship between RAS and Chk1 activity, demonstrating that expression of *KRAS^G12D^* or *HRAS^G12V^* in murine embryo fibroblasts promotes signaling through the ATR-Chk1 pathway^74^. Additionally, Chk1 inhibition in cells with oncogenic mutations in *RAS* induces genomic instability and DNA damage accumulation. In line with this, our findings demonstrate that BRAF inhibitor-resistant cells are hypersensitive to Chk1 inhibition while showing reactivation of MAPK pathway. Furthermore, simulation of a BRAF inhibitor resistance phenotype by PMA-induced activation of the MAPK pathway enhances Chk1 inhibitor sensitivity in treatment-naïve melanoma cells.

Although several mechanisms involved in sensitivity to Chk1 inhibition have been previously proposed, this phenotype likely results from a multitude of genetic and functional alterations, making the identification of a unified mechanism challenging. Our findings indicate that hyperactivation of the MAPK pathway can predict Chk1 inhibition sensitivity, especially in the context of MAPK inhibitor-resistance. However, caution is warranted in using it as a biomarker for patient selection, given the prevalence of MAPK activation across various tumor types, not solely limited to therapy resistance. Therefore, determining the optimal activation levels with clinical significance remains an important unanswered question. Additionally, we cannot assume that hypersensitivity to Chk1 inhibition will be uniform across all MAPK inhibitor-resistant melanomas with re-activation of the MAPK pathway. Our work substantially contributes to the understanding of the effects of Chk1 inhibition in BRAF inhibitor-resistant cells, providing a robust basis for further investigations into the impact of the molecular alterations involved in both resistance to MAPK inhibitors and sensitivity to Chk1 inhibitors. Further investigations employing extensively characterized paired experimental models and patient tumor samples are imperative to identify the subset of patients that can benefit from Chk1 inhibition and to elucidate alternative mechanisms that may play a role in addition to or independent of MAPK pathway activation.

## METHODS

### Lead contact and Materials Availability

Further information and requests for resources and reagents should be directed to the Lead Contact Patricia Abrão Possik.

### Inhibitors and solvents

BRAF inhibitors PLX4720 and Dabrafenib, MEK inhibitor GSK1120212/trametinib and Chk1 inhibitor AZD7762 were purchased from Selleck Chemicals. Chk1 inhibitor GDC-0575 was provided by Genentech. For *in vitro* experiments, all inhibitors were reconstituted in DMSO to a final concentration of 10 mM. Phenylarsine oxide (PAO) and DMSO were purchased from Sigma-Aldrich. Phorbol-12-myristate-13-acetate (PMA) was purchased from Millipore.

### Cell lines and culture conditions

Melanoma and HEK293FT cell lines were cultured in DMEM supplemented with 10% fetal bovine serum (FBS), 3.7mg/ml NaHCO_3_, 1% penicillin-streptomycin 100x, 1% sodium pyruvate 100x and 1% L-glutamine 200mM. All cell lines were confirmed to be mycoplasma free (MycoAlert Plus kit, Lonza, LT07-710) and their identities were confirmed by STR analysis. All melanoma cell lines are *BRAF^V600E^*-mutant. MAPK inhibitors-resistant melanoma cells were derived as described previously (MÜLLER *et al.*, 2014; KEMPER *et al.*, 2016). D4M.3A murine cell line was established from the Tyr::CreER;BrafCA;Ptenlox/lox conditional mouse model (Jenkins, 2014). D4M.3A.BR and D4M.3A.BMR were developed by continuously exposing D4M.3A *in vitro* and *in vivo* to increasing concentrations of Dabrafenib or Dabrafenib plus Trametinib, respectively. Human BRAF inhibitor-resistant melanoma cell lines were maintained in 3μM PLX4720. D4M.3A.BR cells were maintained in 5μM Dabrafenib and D4M.3A.BMR cells were maintained in 1μM Dabrafenib and 0.1μM Trametinib. Inhibitors were removed to perform the experiments.

### Dose Response Curves

Cells were seeded in 96-well plates (1.5-4x10^3^ cells/well) and treated with serial dilutions of PLX4720, Dabrafenib, Trametinib or GDC-0575 for three days. Cell viability was assessed by CellTiter Blue assay (Promega) following the manufacturer’s recommendations. Fluorescence was determined using the SpectraMax ID3 fluorimeter (Molecular Devices) at 560nm and 600nm wavelengths. Values were normalized to vehicle control (DMSO-treated cells) set to 100% viability and killing control (PAO-treated cells) set to 0% viability. Non-linear regression curves were calculated from the average values obtained in three technical replicates using the software Prism GraphPad version 9.0. The half minimal inhibitory concentration (IC50) was estimated for each cell line from at least three independent experiments.

### Apoptosis assay

Cells were seeded in 96-well plates (6x10^3^ cells/well) and treated with GDC-0575 or vehicle control for two days. Images were captured every two hours using Incucyte Caspase-3/7 Green Apoptosis Assay reagent in the Incucyte Live Cell Analysis System (Essen Bioscience). Apoptotic cells were quantified by normalizing green objects to the percentage of cell confluence, based on phase contrast images of each well. Analysis was performed using Incucyte Live Cell Analysis System software. The experiment was performed in at least three biological replicates, each one with three technical replicates.

### Fluorescent Ubiquitination-based Cell Cycle Indicator system (FUCCI)

HEK293FT cells were seeded in 10cm dishes (4x10^6^ cells/dish) and transfected overnight with mKO2-hCdt1 (30/120) or mAG-hGem (1/110) (SAKAUE-SAWANO et al., 2008) and helper plasmids (pMDLglpRRE, pHCMV-G and pRSVrev) using polyethylenimine (PEI) diluted in unsupplemented DMEM medium. Two days after transfection, lentivirus was collected and filtered. For transduction, melanoma cell lines were seeded in six-well plates (3x10^5^ cells/well) and incubated for twenty-four hours with lentivirus carrying the mAG-hGem (1/110) construct. After eight hours of recovery, cells were incubated for twenty-four hours with lentivirus carrying the mKO2-hCdt1 (30/120) construct. Double-positive population was selected by FACS-sorting (FACSAria Fusion SORP cytometer, BD Biosciences; Moflo Astrios, Beckman Coulter; FACSAria IIu, BD Biosciences). After approximately two weeks, cells emitting either green or red fluorescence were FACS-sorted again. This strategy allowed the selection of cells stably expressing both mKO2-hCdt1 (30/120) and mAG-hGem (1/110) that were able to progress through the cell cycle.

### Time-lapse microscopy

FUCCI-labeled cells were seeded in twelve-well plates (15x10^3^ cells/well) and exposed to GDC-0575 or DMSO. Cells were monitored over time using a computer-assisted fluorescence microscope (Zeiss AxioObserver Z1, Zeiss). Images were captured every 20 minutes over 90 hours. The first image was captured approximately one hour after treatment (T_0_). Cell tracking was performed manually in at least sixteen cells in each condition using the Icy software (https://icy.bioimageanalysis.org; DE CHAUMONT et al., 2012). To determine cell cycle duration, untreated cells were tracked from the first frame after cell division until the next cell division. To analyze cell fate, cells were tracked from T_0_ until cell division or the end of the recording period. Red fluorescence (mKO2) was used to determine cells in G1 and green fluorescence (mAG) to determine late S phase, G2 and mitosis. Double labeling was used to determine cells in the G1/S transition and early S phase. The assembly of representative images was performed using the ImageJ software.

### BrdU incorporation

Cells were seeded in 10cm dishes (8x10^5^ cells/dish) and exposed to GDC-0575 or DMSO for six hours. One hour preceding fixation with cold 70% ethanol, cells were incubated with 10µM of BrdU. Samples were stored at -20°C for at least sixteen hours. For double labeling with anti-BrdU and propidium Iodide (PI), samples were blocked with PBS with 1% BSA for five minutes and incubated with a denaturing solution (2N HCl with 0.5% Triton X-100) for thirty minutes. Then, samples were resuspended in 0.1M Sodium Borate solution and washed with PBS with 1% BSA. Labeling was performed with anti-BrdU-FITC mAB (Phoenix Flow Systems, ABFM18) diluted 1:15 in PBS with 1% BSA and 0.5% Tween-20 for one hour. Finally, samples were resuspended in 0.02mg/ml PI with 0.5mg/ml RNAse A diluted in PBS with 0.5% Tween-20 and stored at 4°C in the dark, for three hours. Cell cycle distribution was evaluated by flow cytometry (FACS Calibur) and the frequencies of each phase was estimated by FlowJo software. Analyzes were performed in at least three independent experiments.

### Immunoblotting and antibodies

Cells were seeded in 10cm dishes (1-3x10^6^ cells/dish) and at the end of each experiment, harvested in 1ml of cold PBS. Cell pellets were lysed with RIPA buffer (5 M NaCl; 100% NP40; 20% SDS; 1 M Tris pH 8 and 10% sodium deoxycholate, diluted in Milli-Q water) with protease inhibitors (Protease Inhibitor Cocktail; Sigma, P8340) and phosphatase inhibitors (Phosphatase Inhibitor Cocktail 2; Sigma, P5726; Phosphatase Inhibitor Cocktail 3; Sigma, P0044) by incubation on ice for 30 minutes. Protein concentration was determined by Bradford colorimetric assay (Bio-Rad). Quantified protein lysates were diluted in 5x Laemmli buffer (4.2% Tris; 10% Glycerol; 0.013% Bromophenol Blue 0.1%; 20% SDS; pH 6.8), boiled for 10 minutes and stored at -20 °C until use. Western blotting was performed using 4-20% Mini-PROTEAN TGX gradient gel and the Mini-PROTEAN Tetra Cell system (Bio-Rad). Proteins were transferred to a nitrocellulose membrane (GE Lifesciences) using Mini Trans-Blot module core system (Bio-Rad), followed by blocking with 5% milk powder diluted in TBS-T 0.05% (0.12% Tris; 0.9% NaCl; 0.05% Tween-20) and overnight incubation with primary antibodies diluted in TBS-T with 4% BSA: phospho-RPA32 Ser4/Ser8 (1:1000, Bethyl, A300-245A); үH2AX Ser139 (1:1000, Cell Signaling, 9718); Vinculin (1:2000, Sigma Aldrich, 4650S); Tubulin (1:1000, Sigma, T9026); ERK (1:1000, Cell Signaling, brdu); and phosphor-ERK Thr202/Tyr204 (1:1000, Cell Signaling, 9106). The following secondary antibodies were used: anti-rabbit IgG (1:10,000 -Cytiva, NA934) and anti-mouse IgG (1:10,000 - Thermo Fisher Scientific, 31430), both diluted in 5% milk in TBS-T. Immunoreactivity was detected with ECL SuperSignal West Pico Plus Chemiluminescent Substrate (1:1, Thermo Fisher Scientific) and visualized using a Chemidoc Imaging System (BIORAD). At least two independent experiments were performed.

### DNA fibers Assay

Cells were seeded in six well plates (1.5x10^5^ cells/well) and exposed to GDC-0575 or DMSO for twenty-four hours. Then, cells were incubated with 25μM of 5-chloro-2-deoxyuridine (CldU, Sigma, C6891) for thirty minutes and with 250μM of 5-iodo-2-deoxyuridine (IdU, Sigma, I7125) for thirty minutes at 37°C with 5% CO_2_. Cells were harvested and diluted to 5x10^5^ cells/ml. To spread DNA fibers, 2μl of cells were dropped to the end of the slide (Thermo Fisher Scientific, MNJ-200-010H), incubated for five minutes and mixed with 7μl of the spreading buffer (200 mM Tris-HCl pH 7.4; 50 mM EDTA; 0.5% SDS). After two minutes, the slide was tilted at a 45° angle until the drop reached the other end. The slides were incubated at room temperature for one hour and fixed with 3:1 methanol and acetic acid for ten minutes. After twenty minutes air-drying, the slides were stored at 4°C. For staining, the slides were denatured with 2.5M HCl for seventy-five minutes and blocked with PBS with 1% BSA/0.1% Tween-20 for fifty minutes, followed by one hour incubation with primary antibodies: rat-anti-BrdU (1:500 -Bioconnect clone BU1/75 [ICR1] Jacobslab) and mouse-anti-BrdU (1:750 - Clone B44, Becton Dickinson, 347580 [7580]), both diluted in the blocking solution. After incubation, slides were washed with PBS and fixed with 4% paraformaldehyde for ten minutes. The following secondary antibodies were used: anti-Rat AlexaFluor 555 (1:500 - Invitrogen, A-21434) and anti-Mouse AlexaFluor 488 (1:500 - Thermo Fisher Scientific, A11001), for ninety minutes and diluted in the blocking solution. Two drops of Vectashield mounting medium (Vector laboratories) were added and the coverslip was placed over the slide containing the DNA for thirty minutes in the dark. Images were captured using Zeiss AxioObserver Z1 inverted microscope equipped with a black and white Hamamatsu ORCA AG CCD refrigerated camera at 63x magnification. The length of the DNA fibers was measured using the ImageJ software. Replication forks progression speed was quantified by the length of the DNA strands labeled with CldU or IdU and converted to kilobases (kb) incorporated per minute (kb/min). Filaments with features of progressing forks, origin firing and termination were quantified. Experiments were performed in two biological replicates.

### RNA-sequencing analysis

Cells were seeded in 10cm dishes (1x10^6^ cells/dish) and exposed to 0.3µM of GDC-0575 for six hours. As a control, cells were harvested before treatment (T_0_). Experiments were performed in three independent biological replicates. RNA was extracted using the RNeasy Plus Mini kit (Qiagen) and quantified with Qubit RNA Broad Range kit and the Qubit 4 (Invitrogen). RNA quality was assessed using the RNA 6000 Nano kit and the 2100 Bioanalyzer (Agilent Technologies). Library was prepared with TruSeq Stranded mRNA kit (Illumina). Samples were indexed and analyzed by paired-end sequencing of 75 base pairs on the HiSeq 2500 instrument (Illumina). Sequenced reads were trimmed to remove low-quality bases using Trimmomatic v0.39 (SLIDINGWINDOW:4:20 LEADING:3 TRAILING:3 MINLEN:36)^75^. The default parameters of STAR program version 2.6.0c (https://github.com/alexdobin/STAR/releases;^76^) was used to map the clean reads using GRCh38 genome (Ensembl) as reference and to quantify the number of reads per gene, using the *"*quantMode GeneCounts*"* option. To detect differentially expressed genes, the reads matrix was normalized using the R software version 4.0.5 (http://www.R-project.org/; R CORE TEAM, 2013), with DESeq2 package version 1.28.1^77^. A pre-filtering was performed to exclude genes with less than 10 reads in all samples. To estimate the log2 fold change, DESeq2 package was used to adjust the p value from Wald Test using Benjamin and Hochberg method. Only the resistant cells were used in the design. For this analysis, four comparisons were performed: 888melR control vs 888mel control; 888melR treated with

GDC-0575 vs 888mel treated with GDC-0575; 888melR treated with GDC-0575 vs 888melR control; 888mel treated with GDC-0575 vs 888mel control. Genes with p-adjusted ≤ 0.05 were considered differentially expressed (Wald Test). Gene set enrichment analysis (GSEA) was performed with clusterProfiler package version 3.16.1^78^, using KEGG pathway database (https://www.genome.jp/kegg/kegg1.html;^79–81^). Heatmaps were generated using the pheatmap package (RRID:SCR_016418). Box plots were made using the ggplot2 package. The expression data of a set of MAPK-specific genes (*PHLDA1*, *SPRY2*, *SPRY4*, *DUSP4*, *DUSP6*, *CCND1*, *EPHA2*, *EPHA4*, *ETV4*, and *ETV5*) was used to calculate the MAPK Pathway Activity Score (MPAS). Normalized RNAseq read counts were used to calculate the Z-score of each gene’s expression. MPAS was computed accordingly to the following formula, where *z_i_* is the z-score of each gene’s expression level, and *n* is the number of genes comprising in the set (i.e., n = 10)^82^: All analyzes were performed in the R software.

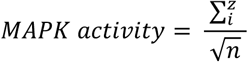

### Animal experiment

Cells were prepared in a 1:1 mixture of growth factors-reduced matrigel (Corning, 354230) and supplemented DMEM medium and inoculated subcutaneously into NOD/SCID IL2Rgamma null (NSG) immunodeficient mice (1x10^6^ cells/mouse). On the first day after inoculation, mice were randomly allocated into two groups (n=4 mice each) and treated with 25mg/kg of AZD7762 or vehicle (11.3% 2-hydroxypropyl-β-cyclodextrin in 0.9% NaCl (pH 4) -HP-β-CD). Treatment was administered intraperitoneally three times a week. Tumor volume was monitored three times a week and calculated using the formula ½ x length (mm) x width (mm). The experimental endpoint was 1000 mm^3^ of tumor volume in at least one animal in the group, which occurred on the 21^st^ day of treatment for the control groups. None of the mice treated with AZD7762 reached 1000 mm^3^ of tumor volume and the experiment ended on the 31^st^ day of treatment. The experiment was carried out according to the local and international regulations and ethical recommendations. The experiment was authorized by the animal experimentation committee of the Netherlands Cancer Institute (protocol number 1.1.9231).

### Statistical testing

*In vivo* tumor growth data was analyzed at the 31^st^ day of treatment by unpaired Student’s t-test. For *in vitro* experiments, either the Student’s T test or the Mann-Whitney test was employed to compare two conditions, depending on whether the data were normally distributed. For comparing two or more conditions, unpaired one-way ANOVA or the Kruskal-Wallis test was performed, depending on whether or not a normal distribution was observed. Normality was assessed by the Shapiro-Wilk test or D’Agostino & Pearson test. For ANOVA, the Tukey post-test was performed to compare all experimental groups. For Kruskal-Wallis, the Dunn post-test was conducted to correct for multiple comparisons. Curves obtained through caspase 3/7-mediated cleavage were analyzed using the Friedman test. To evaluate if there is an association between two factors, Chi-square and Fisher’s exact test was performed. The analyses were carried out using Prism GraphPad version 9.0, which was also used for generating curves and graphs. Values with a null hypothesis probability (p) equal to or less than 0.05 were considered significantly different.

## Data availability

The RNA sequencing dataset related to this study and the code used to analyze it will be deposited in relevant public databases upon acceptance of this article for publication.

## AUTHOR CONTRIBUTIONS

D.S.P and P.A.P conceived and supervised the study. D.G.C. and J.C.N.K. performed all experiments with support from M. A. R and C. F. O. K, D. A. M and M. B performed the bioinformatics analyses. J.P.B.V provided critical input. D.G.C., J.C.N.K and P.A.P wrote the manuscript. All authors reviewed and approved the manuscript.

## ACKNOWLEDGEMENTS

We would like to thank all members of Possik and Peeper laboratories for helpful advice throughout this project. Special thanks to Daniel Postrach from German Cancer Research Center (DKFZ) and David J. Adams, from Wellcome Sanger Institute for support, discussions and reading of the manuscript. We would also like to thank the members of The NKI’s Bioimaging Facility, Karina Lani of the Cytometry Core Facility at INCA, Nicole Scherer of the Bioinformatics Core Facility at INCA, and the Microscopy and Bioimaging Facility at INCA. This work was financially supported by Science Without Borders Program-CAPES (J.C.N.K., grant #1334713-6), Science Without Borders Program-CNPq (P.A.P, grant # 401116/2014-0), International Centre for Genetic Engineering and Biotechnology - CRP-ICGEB Early Return Grant (P.A.P., grant # CRP/BRA17-05_EC) and Fundação de Amparo a Pesquisa do Estado do Rio de Janeiro (P.A.P., grants # E-26/010.001645/2019 and E26/010.002187/2019). D.G.C. was supported by a fellowship from INCA and the Ministry of Health.

## SUPPLEMENTAL FIGURE TITLES AND LEGENDS

**Supplementary Figure 1, related to Figure 1: Sensitivity to Chk1 inhibition is exacerbated in cells resistant to BRAF and MEK inhibitors.** A) Treatment-naïve parental (P), BRAF inhibitor-resistant (BR) and BRAF and MEK inhibitors-resistant (BMR) D4M.3A mouse melanoma cells were treated with different concentrations of GDC-0575 for 72 hours. Cell viability was determined by CellTiter Blue. Percentage of viable cells are depicted as mean (+/- SD). B) IC50 values were estimated from the curves (A) and represented as mean (+/- SEM). n = 5 biological replicates, with 3 technical replicates each. Ordinary one-way ANOVA and Tukey’s multiple comparisons test. ** p < 0.01.

**Supplementary Figure 2, related to Figure 1: Sensitivity to Chk1 inhibition is a consequence of increased apoptotic cell death.** A) Treatment-naïve parental (P) and BRAF inhibitor-resistant (R) melanoma cells were exposed to GDC-0575 or DMSO for 48 hours and monitored using time-lapse microscopy, as described and represented at Figure 1B. The mean of each biological replicate at 48 hours of GDC-0575 treatment (+/- SD) was normalized by control samples at the same time-point. n = 3 biological replicates with 3 technical replicates each. Two-way ANOVA. **** p < 0.0001. B) 888mel and 888melR cells were inoculated s.c. into NSG mice. Vehicle or AZD7762 (25mg/kg) treatment was performed 3x week until mice reached 800mm^3^. n=4 mice per group. Mantel-Cox Test. ** p < 0.01.

**Supplementary Figure 3, related to Figure 2: BRAF inhibitor-resistant and treatment-naïve melanoma cells show similar duration of the cell cycle.** A) Representative images of FUCCI-labeled cells followed by time-lapse microscopy for up to 90 hours. No treatment was administered. B-C) The duration of G1, early S and S/G2/M phases was determined by tracking 20 parental (P) and BRAF inhibitor-resistant (R) cells according to FUCCI’s fluorescence. Total cell cycle indicates the sum of G1, early S and S/G2/M phase duration times. Unpaired Student’s T test. **** p < 0.0001.

**Supplementary Figure 4, related to Figure 2:** A) Representative images of distinct phenotypes observed in FUCCI-labeled cells followed by time-lapse microscopy (up to 90 hours). B-C) FUCCI-labeled parental (P) and BRAF inhibitor-resistant (R) melanoma cells treated with GDC-0575 (0.3µM for 888mel/888melR and 0.1µM for M026.X1.CL/M026R.X1.CL) and followed by time-lapse microscopy. Cells that failed to complete the cell cycle upon Chk1 inhibition were categorized according to the acquired phenotype (11-24 cells were analyzed). D-E) Cells that acquired a rounded morphology upon Chk1 inhibition were classified according to the cell cycle phase observed at the end of tracking (6-24 cells were analyzed). Statistical significance was calculated using Chi-squared and Fisher’s exact test. p values are indicated.

**Supplementary Figure 5, related to Figure 3: Chk1 inhibition does not consistently modulate cell cycle distribution in BRAF inhibitor-resistant melanoma cells.** A) Treatment-naïve (P) and BRAF inhibitor-resistant (R) cells were treated with GDC-0575 (0.3µM for 888mel/888melR and A375/A375R or 0.1µM for M026.X1.CL/M026R.X1.CL) or DMSO for 6 hours. Cells were pulsed with BrdU for 1 hour and analyzed by flow cytometry. n ≥ 3 biological replicates. Representative graphs are shown. B) Frequency of cells in G1, S and G2/M phases. Ordinary one-way ANOVA or Kruskal-Wallis. * p < 0.05; ** p < 0.01. *** p ≤ 0.001. Only significant comparisons are represented.

**Supplementary Figure 6, related to Figure 3: Chk1 inhibition suppresses DNA replication and repair in both parental and BRAFiR cells.** A) Schematic representation and micrographs of DNA fibers assay from treatment-naïve (P) and BRAF inhibitor-resistant (R) cells upon GDC-0575 treatment (0.3µM for 24 hours). B) Nucleotide incorporation speed rate (kilobase/minute). n = 2 biological replicates, and only one is represented. Ordinary one-way ANOVA. **** p < 0.0001.

**Supplementary Figure 7, related to Figure 4: DNA Damage Response and DNA Synthesis pathways are altered in melanoma cells upon Chk1 inhibition.** RNA sequencing of 888mel and 888melR cells untreated (P0 and R0, respectively) or treated with GDC-0575 for six hours (P6 and R6). Gene Set Enrichment Analysis (GSEA) illustrates main altered pathways. Only pathways with GeneRatio > 0.4 were considered. n = 3 biological replicates.

