## Supplemental files for "Resistance to BRAF inhibitors drives melanoma sensitivity to Chk1 inhibition"

**Supplementary Figure 1**

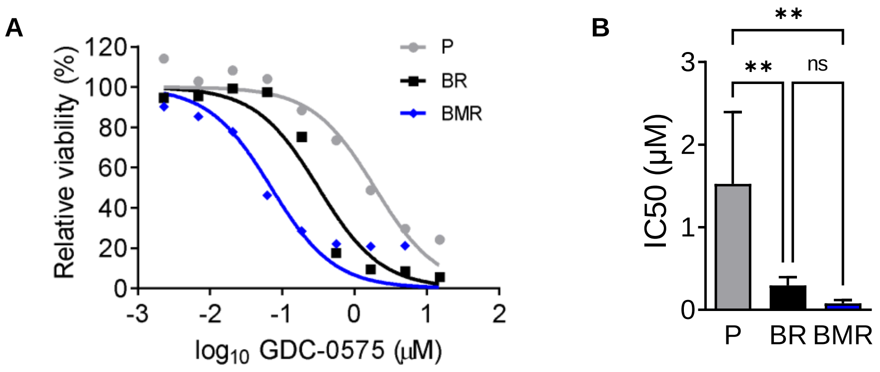

**Supplementary Figure 2**

**A**

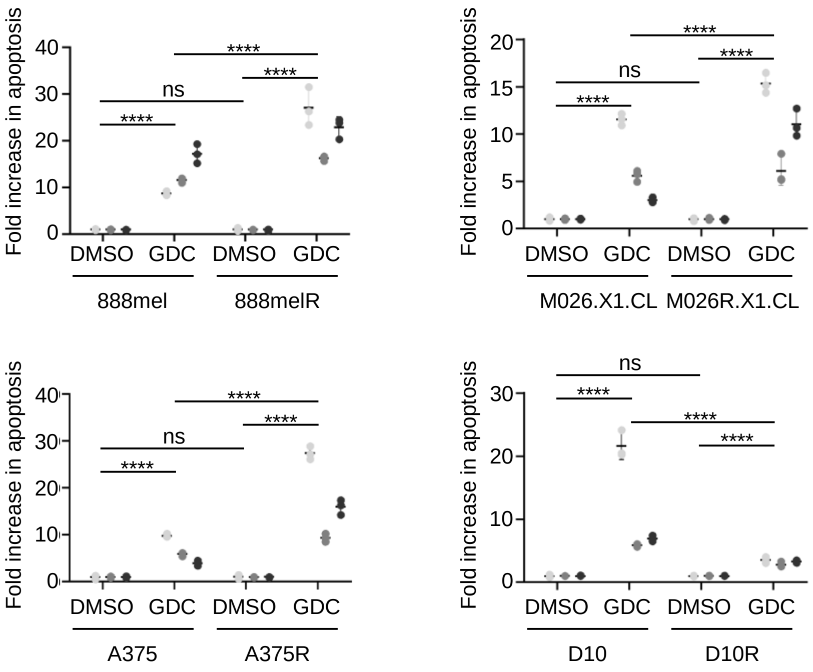


**B**

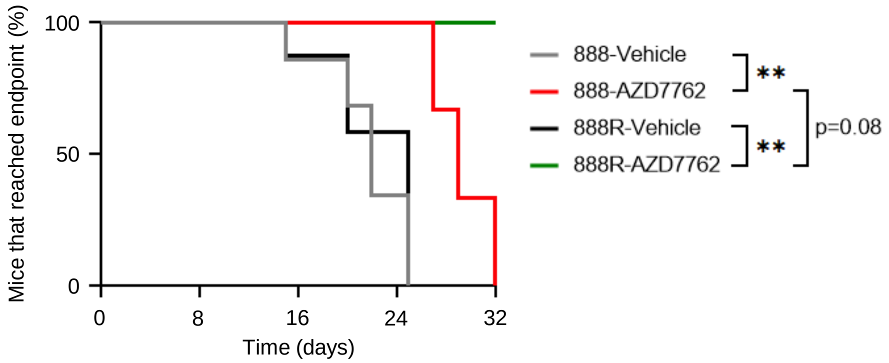

**Supplementary Figure 3**

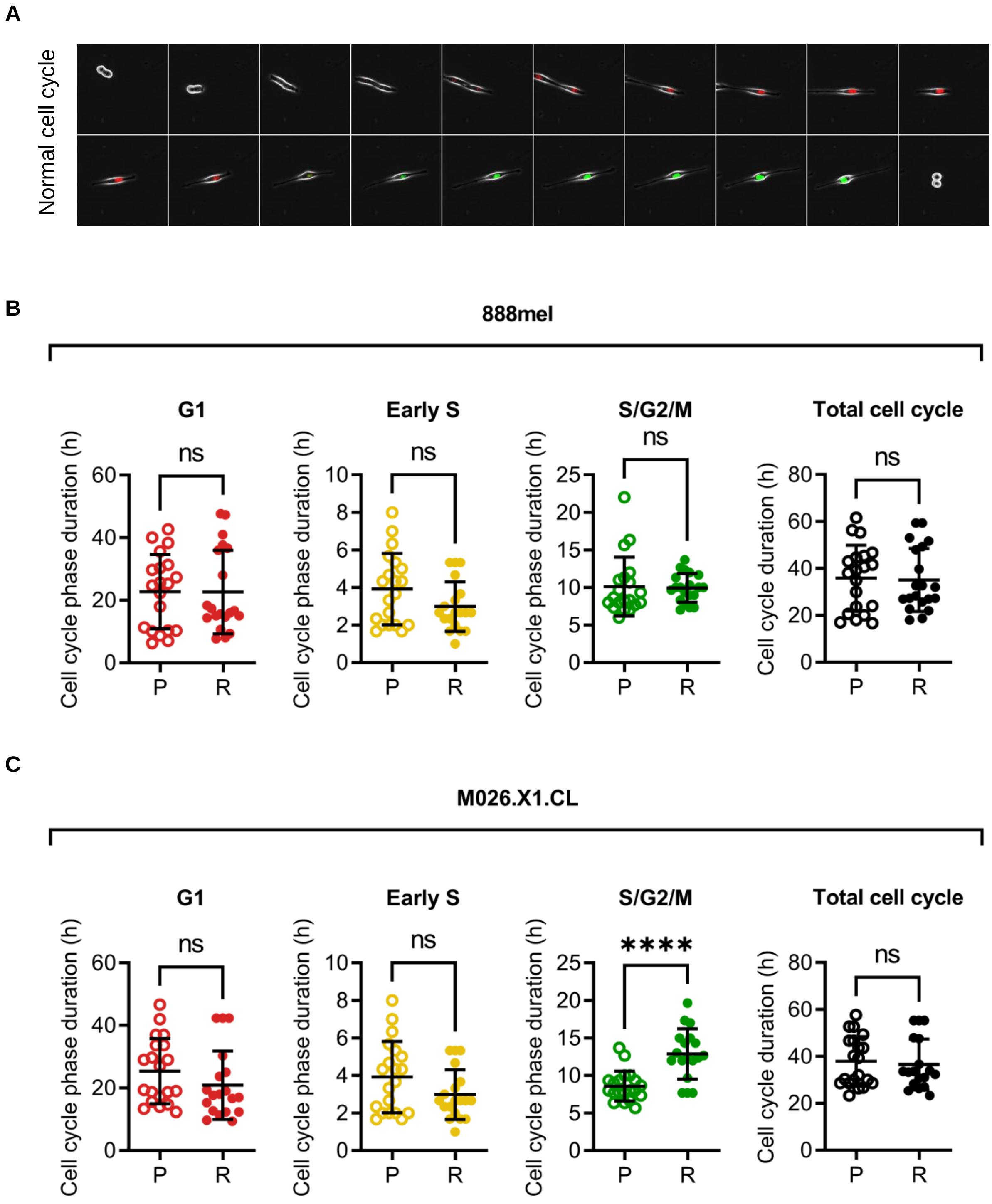

**Supplementary Figure 4**

**A**

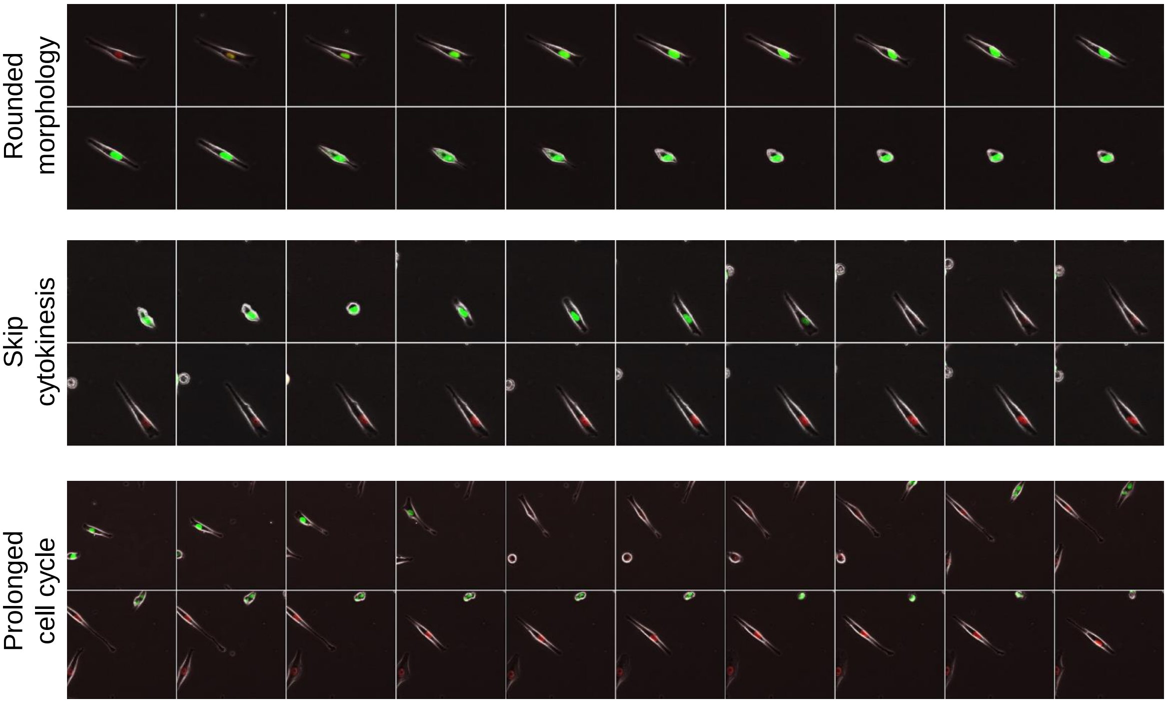


**B**

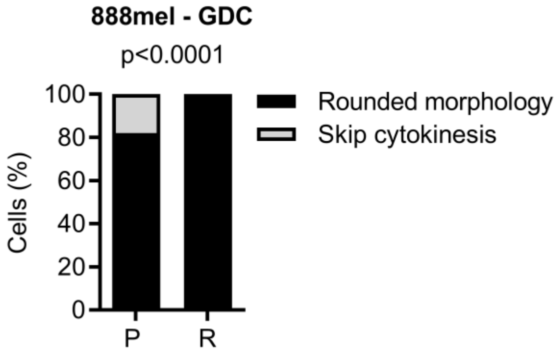


**C**

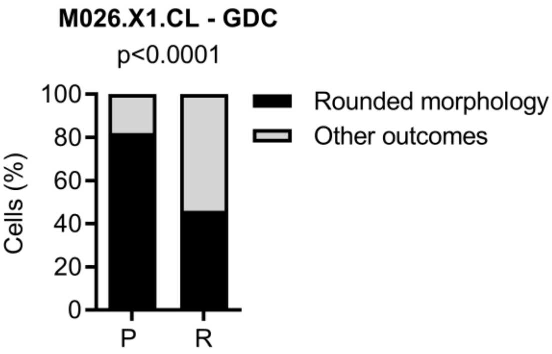


**D**

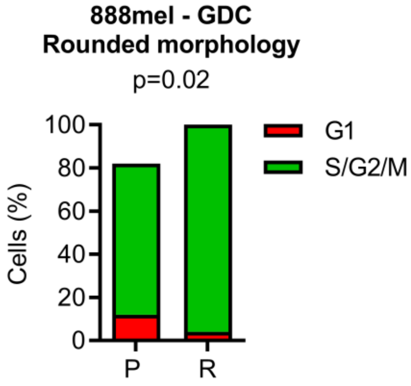


**E**

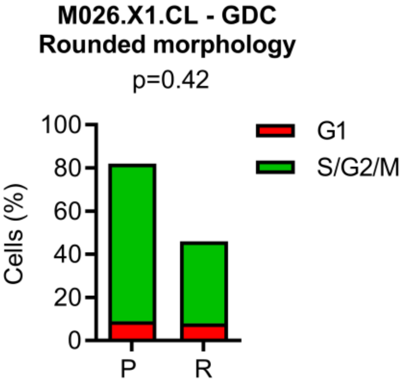

### Supplementary Figure 5

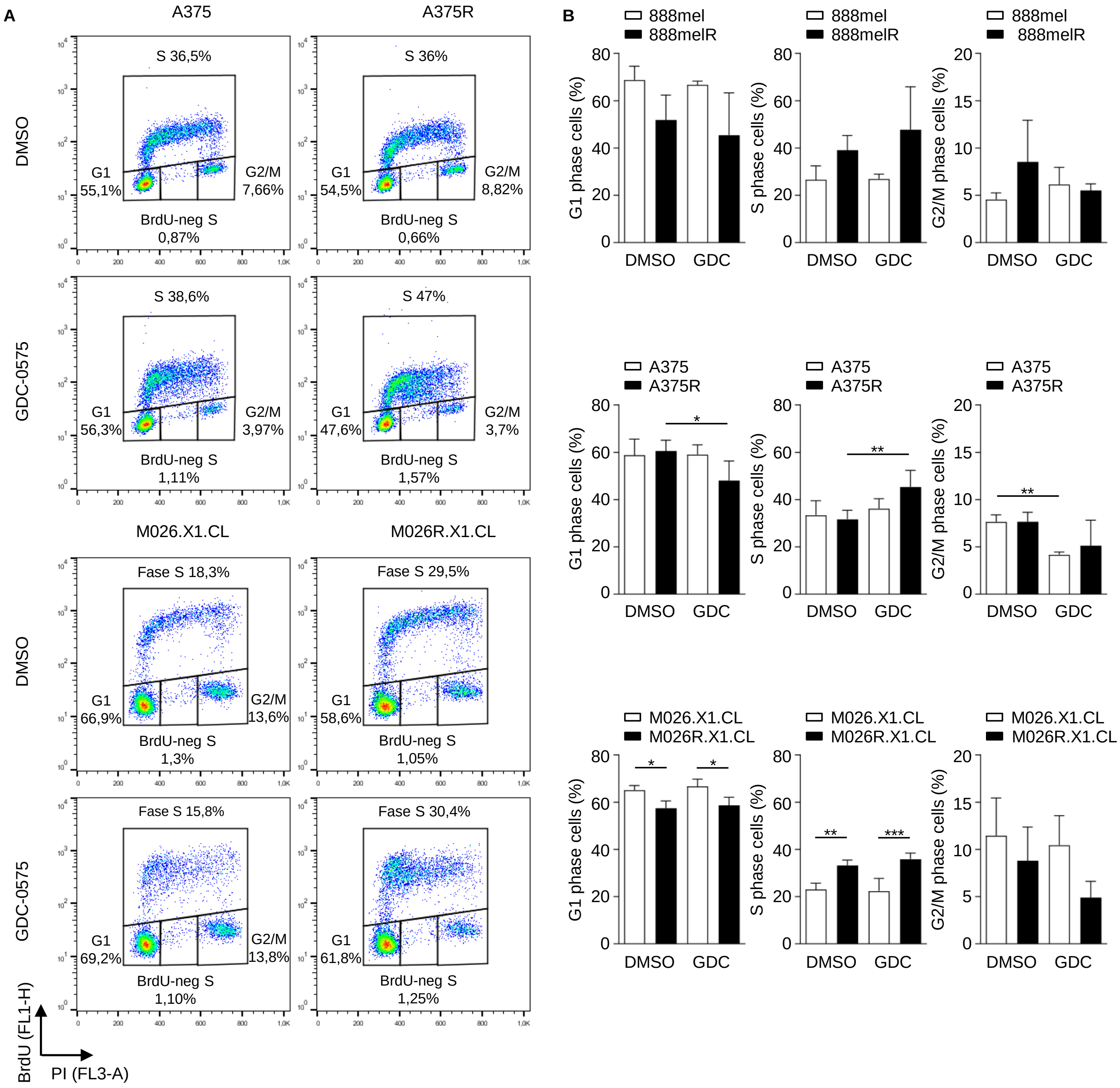

### Supplementary Figure 6

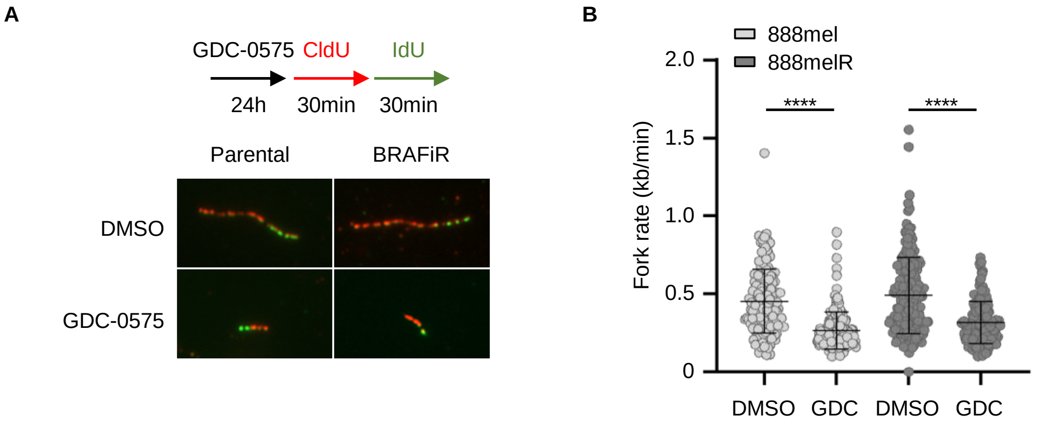

### Supplementary Figure 7

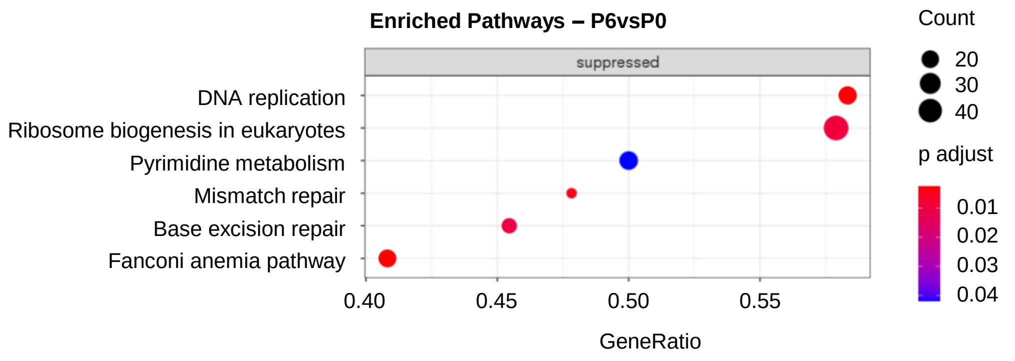


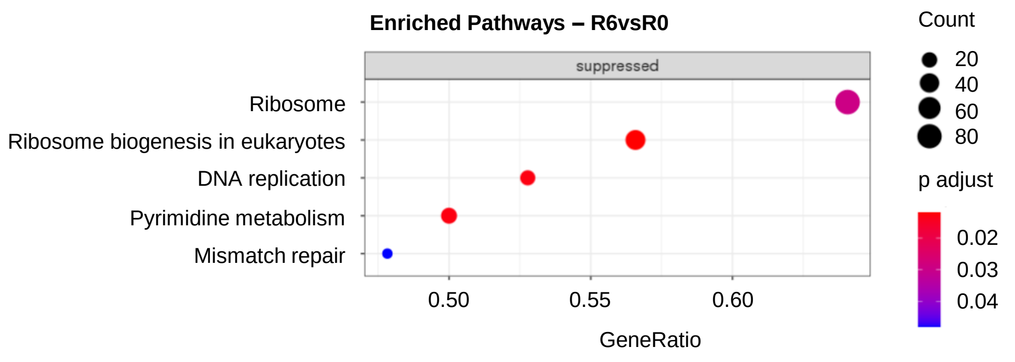
